# “Plasma Membrane Calcium Atpase downregulation in dopaminergic neurons alters cellular physiology and behavior in Drosophila melanogaster”

**DOI:** 10.1101/714147

**Authors:** Brenda Erhardt, María Celeste Leal, María Silvina Marcora, Lía Frenkel, Pablo Alejandro Bochicchio, Diego Hernán Bodin, Berenice Anabel Silva, María Isabel Farías, Carina Cintia Ferrari, Miguel Ángel Allo, Christian Höcht, Eduardo Miguel Castaño, Fernando Juan Pitossi

**Author notes:** Correspondence should be addressed to Dr. Fernando J Pitossi, IIBBA-CONICET, Fundación Instituto Leloir, Av. Patricias Argentinas 435 C1405BWE, CABA, Argentina. /1 (ext 3106). both authors contributed equally to this work (Brenda Erhardt, María Celeste Leal).

## Abstract

Accumulation of calcium is proposed to account for selective dopaminergic neuron (DN) dysfunctionality, a characteristic of Parkinson’s Disease (PD). To test the *in vivo* impact of calcium increment in DN physiology we downregulated the Plasma Membrane Calcium ATPase (PMCA), a bomb that extrudes cytosolic calcium, in those neurons in *Drosophila melanogaster.* Using *th*-GAL4>PMCA^RNAi^, PMCA was selectively reduced, leading to increased cytosolic calcium and mitochondrial oxidative stress with no neurodegeneration. In the eye, PMCA^RNAi^ expression provoked a subtle disorganization, suggesting scarce toxicity. Interestingly, we observed several locomotor alterations and a higher level of dopamine in brains. Finally, flies presented a reduction of lifespan and a *perimortem* non-motor phenotype characterized by abdominal swelling, possibly due to constipation. We conclude that elevated cytosolic calcium in DN could trigger cellular dysfunction generating mitochondrial oxidative stress and motor and non-motor symptoms, typical of PD.

## Introduction

Dysfunction of dopaminergic neurons (DN) underlies motor and non-motor symptoms in Parkinson’s disease (PD). Signs of PD are tremor, trouble moving or walking, constipation, among others [1]. Why DN in *substantia nigra* (SN) are particularly vulnerable is one of the key unanswered questions in PD research. Particular combination of unique features of DN-broad spikes, pacemaker activity, low intrinsic calcium (Ca^2+^) buffering and cytosolic Ca^2+^ oscillations-has been proposed to increase DN vulnerability. The pacemaker firing of DN is dependent on Ca^2+^ influx. It has been hypothesized that such elevated Ca^2+^ influx is associated with a considerable metabolic cost and reactive oxygen stress (ROS) increment [2]. The elevated rate of mitochondrial oxidative phosphorylation required to meet the cell’s demands in ATP has been hypothesized to lead to oxidative stress and contribute to the heightened vulnerability of DN [3].

*Drosophila melanogaster* (DM) has yielded great advances into the underpinnings of many neurological and neurodegenerative diseases, not only providing understanding of biological pathways impaired in disease, but also the foundation for intervention strategies in mammalian systems. The approach of recreating a simplified version of human neurological disease by expressing (or knocking down) a human mutant gene in the fly has now been used to model features of many neurodegenerative disorders [4]. To test *in vivo* the relevance of cytosolic Ca^2+^ increment in DN we downregulated Plasma Membrane Ca^2+^ ATPase (PMCA) expression in DM. As PMCA is a bomb that extrudes Ca^2+^, the reduction of its expression using *th-*GAL4 to drive PMCA^RNAi^ in DN could be a good model to increment cytosolic Ca^2+^.

## Results and discussion

### 1-Downregulation of PMCA on DN causes Ca^2+^ and ROS increments

To analyze the relevance of PMCA *in vivo*, we used an RNAi to downregulate PMCA expression in DM. Expression of PMCA^RNAi^ using the panneural promoter *elav* produced unviability (*elav*-GAL4>PMCA^RNAi^, data not shown). We then, selectively expressed PMCA^RNAi^ in DN using a *th* driver. We obtained a ∼40% of reduction in PMCA expression in the head of these flies studied by qRT-PCR (Figure S1A). We then measured Ca^2+^ levels in DNs using a GCaMP reporter. As somas are in different planes we measured GCaMP fluorescence in a single plane in axons net, as a representation of the Ca^2+^ levels in several DN clusters. Fluorescence was quantified in *ex vivo* brains from flies *th-*GAL4>GCaMP/PMCA^RNAi^ and *th-*GAL4>GCaMP/ChRFP as control (Figure 1B vs A). ChRFP signal as control is in Figure S1B. DN axons expressing PMCA^RNAi^ showed more fluorescence than control, indicating that the expression of PMCA^RNAi^ in DNs generated higher levels of Ca^2+^ in dopaminergic projections (Figure 1C).

**Figure 1.**
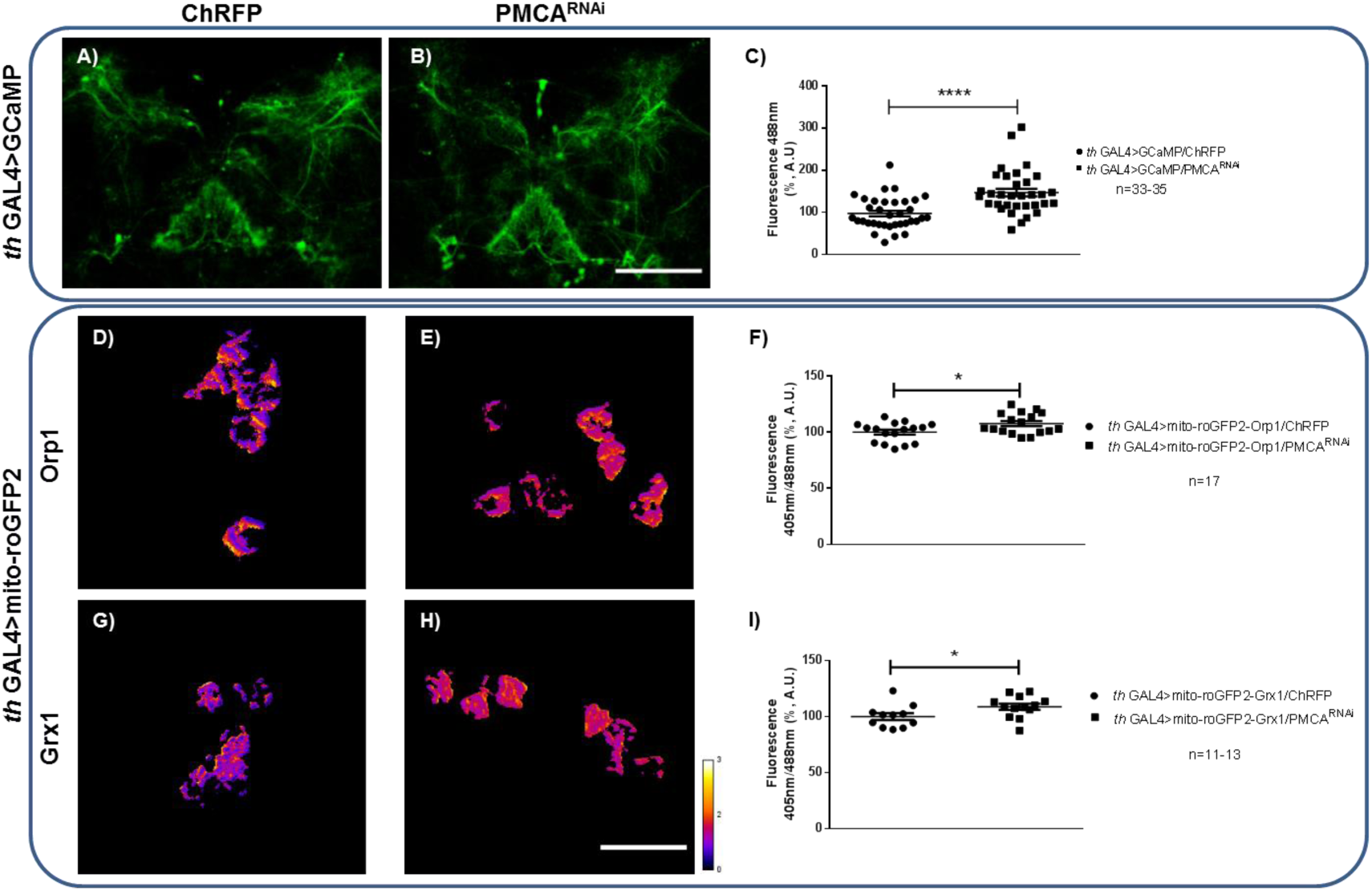
PMCA knockdown in DNs produced basal Ca^2+^ and ROS levels raises. A-C, Ca^2+^ basal state, measured by GCaMP fluorescence in DN axons. Representative images of DN axons co-expressing GCaMP and A, ChRFP or B. PMCA^RNAi^. C, GCaMP fluorescence measurement showed Ca^2+^ increments in DN expressing PMCA^RNAi^. D-I, ROS measured by mitochondrial roGFP2 probes. Representative images of the ratio 405nm/488nm in false color scale from PPM3 somas co-expressing mito-roGFP2-Orpl and D. ChRFP or E. PMCA^RNAi^ and mito-roGFP2-Grxl and G. ChRFP or H. PMCA^RNAi^. Quantification of 405nm/488nm fluorescence ratio showed a significant increment in mitochondrial H_2_O_2_ (F) and GSSG levels (I) in PMCA^RNAi^ expressing PPM3. A, B. Scale bar: 100 μn. D, E, G, H. Scale bar: 15 pm. t test *p<0.05, ****p<0.0001.

To explore if the increment in Ca^2+^ levels in DN produced an increment in ROS levels, we used mitochondrial and cytosolic roGFP2 fused to Orp1 and Grx1 as reporters of H_2_O_2_ and glutathione disulfide (GSSG), respectively [5]. We measured ROS levels in PPM3 DN cluster somas, as they are functionally homologous to those of the mammalian SN [6]. We found that H_2_O_2_ and GSSG are increased in mitochondria of PPM3 somas expressing PMCA^RNAi^ compared to control (PPM3 somas expressing ChRFP), denoted by roGFP2-Orp1/roGFP2-Grx1 reporters (shown as the ratio of 405nm/488nm in false-color images, fire scale) (Figure 1D-I). It has been demonstrated that Ca^2+^ influx during pacemaker activity increases mito-roGFP2 in DN from SN in culture, being prevented by isradipine Ca^2+^ blocker [7].

On the other hand, we measured cytosolic ROS levels with cyto-Grx1-roGFP2 in PPM3 somas. There was no difference between somas expressing PMCA^RNAi^ compared to control (Figure S1C-E). No signal was detected of cyto-Orp1-roGFP2 reporter (data not shown). The observed differences between cytosolic and mitochondrial GSH-redox potential are compatible with the notion that cytosolic and mitochondrial GSH pools are independently regulated [8]. Measurement of mito-roGFP2-Orp1 reporter in the axons net did not show differences of H_2_O_2_ levels (Figure S1F-H) and mito-roGFP2-Grx1 and cyto-Grx1-roGFP2 reporters were not measurable due to low 405nm signal and trachea autofluorescence (data not shown). We conclude that a reduction in PMCA expression in DN lead to increased Ca^2+^ and ROS levels, a feature of many models of PD [2, 7, 9-10].

### 2-PMCA^RNAi^ expression causes scarse toxicity and no neurodegeneration

To analyze if Ca^2+^ and ROS elevated levels affect cellular viability, we quantified DN number by immunohistochemistry using anti-tyrosine hydroxylase (TH) staining as marker. We found no differences between the number of TH positive cells in DN PPL1, PPL2, PPM1/2 and PPM3 clusters of flies expressing PMCA^RNAi^ and controls. This result implied that there is no overt DN death (Figure 2A).

**Figure 2.**
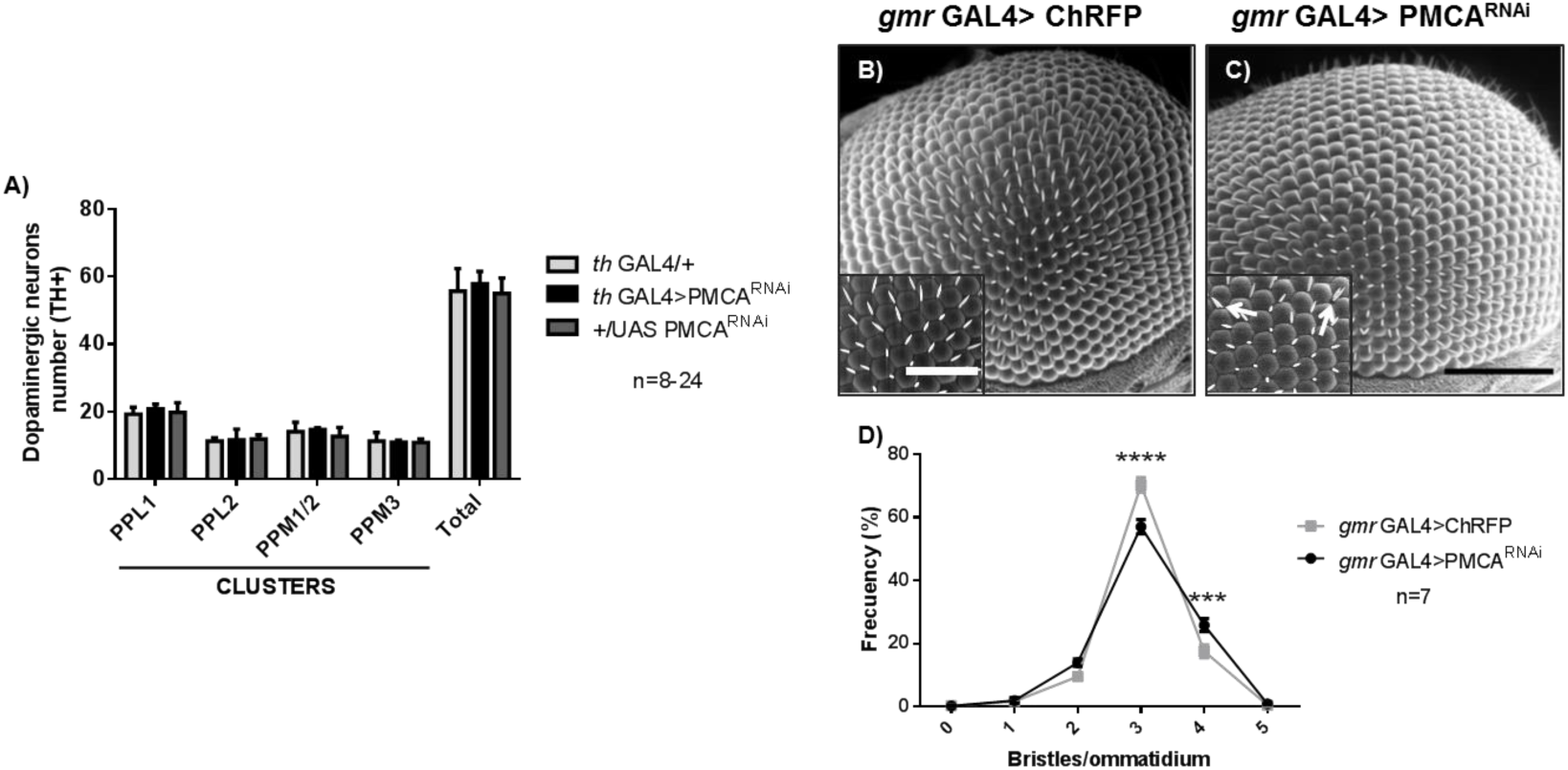
PMfCA^RNAi^ expression did not induce cellular death and generated subtle toxicity in eye context. A, Number of DN TH+ for PPL1, PPL2, PPM 1/2 and PPM3 clusters measured by immunostaining. No differences were observed. B-D, Morphological analysis by SEM of eye integrity (*gmr* driver). Representative images of eyes expressing B, ChRFP or C, PMCA^RNAi^. In the inset panel, a higher magnification shows duplication of bristles (indicated with arrows). D, Quantification of frequency of the number of bristles per ommatidium for each genotype. Eyes expressing PMCA^RNAi^ presented a higher frequency of 4 and lower of 3 bristles per ommatidium. B, C Scale bar: 100 μm, inset panel: 50 μm. A, Two-way ANOVA, Tukey test; n.s. p>0,05. D, Two-way ANOVA, Sidak test; ***p<0,001, ****p<0.0001.

Then, we expressed PMCA^RNAi^ in the eye, to visualize possible, more subtle toxic effects, using *gmr* driver. Each ommatidium has 3 bristle complexes in an organized structure [11]. Eyes expressing PMCA^RNAi^ or ChRFP as control were subjected to scanning electron microscopy (Figure 2C and 2B, respectively). We found a subtle disorganization phenotype proven by the quantification of the number of bristles per ommatidium. *gmr*-GAL4>PMCA^RNAi^ eyes showed significantly lower ommatidia with 3 and more with 4 bristles than control eyes (Figure 2D). These results suggest that reduction of PMCA through PMCA^RNAi^ provoke scarse cellular toxicity at the morphological level.

### 3-Expression of PMCA^RNAi^ in DN produces alterations in locomotor activity and an increment in dopamine levels

To correlate cellular phenotypes with a behavioral response, we analyzed locomotor activity. We performed climbing assays [12]. Climbing ability has been shown to be sensitive to oxidative stress [13]. We found that *th*-GAL4>PMCA^RNAi^ flies climbed less than controls at 10 day after eclosion (Figure 3A). No difference was observed in previous times. To evaluate the general walking status of the PMCA^RNAi^ expressing flies we additionally used the open field assay similar to [14]. We measured walking parameters including total distance walked and maximum time of detention. We found a trend that PMCA^RNAi^ expressing flies walk less than controls when measured the total distance tracked (Figure S2A-B), although differences were not statistically significant among all groups. When we analyzed the time of detention, we found that *th*-GAL4>PMCA^RNAi^ flies remain stopped more time than control genotypes, at 2 and 10 days post-eclosion (Figure 3B-C). Since PMCA reduction caused flies to stand immobile, we speculate that this phenotype could simulate motor dysfunctions and a *freezing* condition similar to freezing of gait observed in PD patients [15].

**Figure 3.**
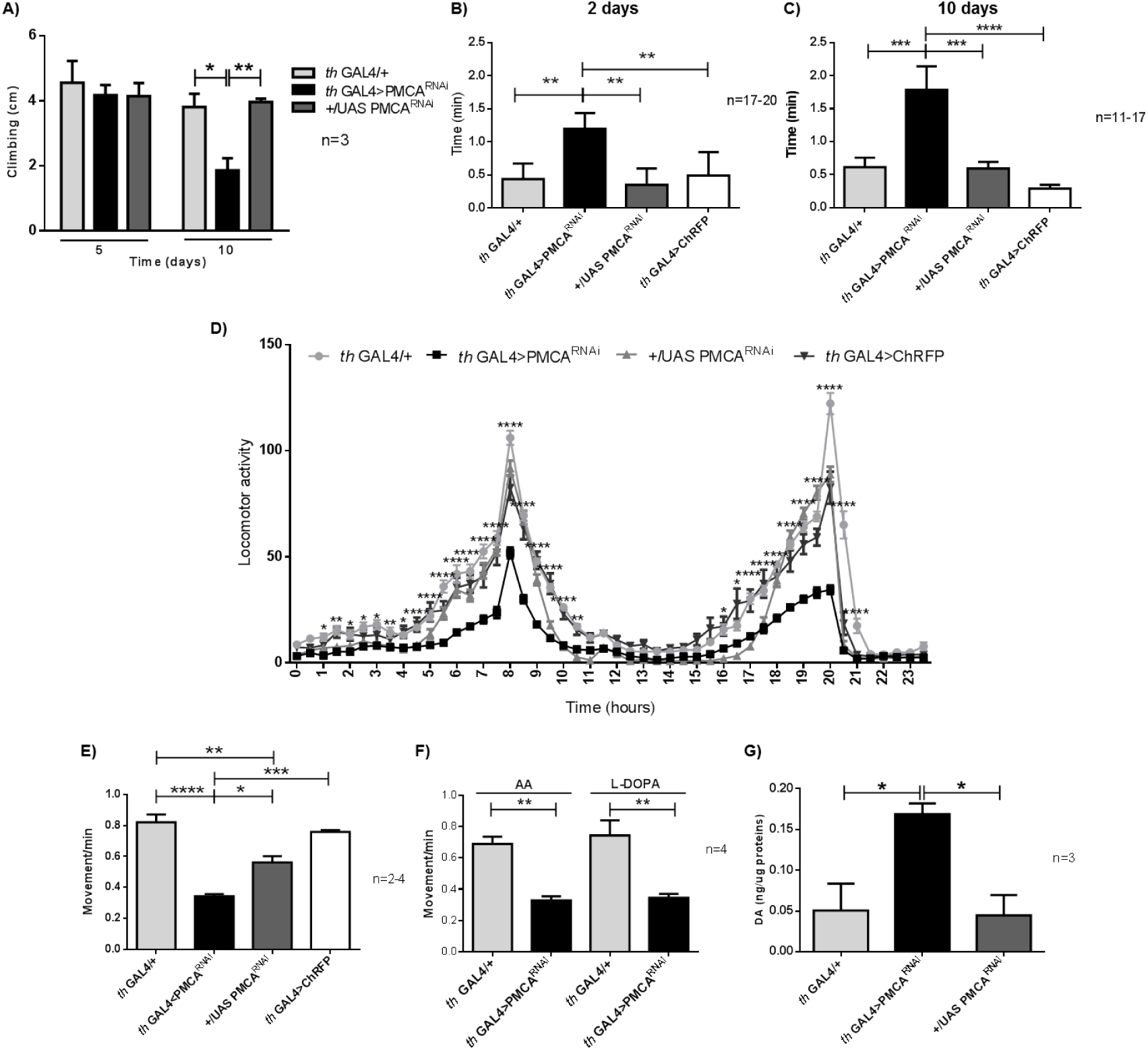
Locomotor activity was altered and dopamine levels incremented in flies expressing PMCA^RNAi^ in DN. A, Climbing ability. Flies expressing PMCA^RNAi^ showed climbing alterations at 10 days after eclosion, compared to controls. B-C, Time of detention. Flies expressing PMCA^RNAi^ were still two (2 days, B) or three (10 days, C) times more than other genotypes. D-F, Monitoring of total locomotor activity. D, Mean temporal organization of locomotion showed lower movement from PMCA^RNAi^ flies (black) vs controls (grays) during the peaks of activity. E-F, Quantification of total locomotor activity. Flies expressing PMCA^RNAi^ moved less than controls (E) and this lower total activity did not revert with L-DOPA addition (F). G, Dopamine levels measured by HPLC. Brains from PMCA^RNAi^ expressing flies had more dopamine than controls. A, Two-Way ANOVA, Tukey test; B-G, One-Way ANOVA, Tukey test; *p<0.05, **p<0.01, ***p<0.001, ****p<0.0001.

As a third independent technique we used a locomotor activity fly paradigm. Total activity was monitored and data was collected every minute per 10 days. In Figure 3D we show a histogram of activity from *th-*GAL4>PMCA^RNAi^ vs. control genotypes. Flies expressing PMCA^RNAi^ presented less activity as compared to control flies, easily observable at times when flies show major peaks of activity (∼ 7:30 am or pm). Graph in Figure 3E shows total activity accumulated. *th*-GAL4>PMCA^RNAi^ flies moved less than controls during their whole life, seen at 3-5 and 8-10 days (Figure S2C-D). We hypothesized that the physiological changes in DN are responsible for performance deficits observed in all these assays. Future experiments of detailed cellular physiology are necessary to explain locomotor alterations. To determine the specificity of results explained above, we included L-DOPA in fly food, in an attempt to revert locomotor alterations. As shown in Figure 3F, exogenous L-DOPA did not revert the reduction in total activity. In sight of these results, we measured dopamine levels in flies’ brains by HPLC. *th*-GAL4>PMCA^RNAi^ flies had higher dopamine levels than controls (*th*-GAL4/+ 0.051±0.033; *th*-GAL4>PMCA^RNAi^ 0,169±0.013; +/UAS-PMCA^RNAi^ 0.045±0.025, ng dopamine/µg total protein; Figure 3G). Locomotor deficits and dopamine increment without DN loss are similar to findings observed in parkin KO mice and flies [16, 17]. These results prompt us to hypothesize that the life span of DM is too short to detect neurodegeneration or TH staining is not sensible enough to detect changes but its lifespan is sufficient to allow the observation of functional deficits as seen in previous DM models [18, 19].

### 4- th-GAL4>PMCA^RNAi^ have shorter lifespan by constipation

We analyzed survival curves, and we found that *th*-GAL4>PMCA^RNAi^ flies died before controls, with a median survival of 13 days (Figure 4A). In a simple empirical analysis, flies *th-*GAL4>PMCA^RNAi^ showed a *perimortem* swelling phenotype in abdominal area (it appears just ∼10 days post eclosion), compared to controls (Figure 4B).

**Figure 4.**
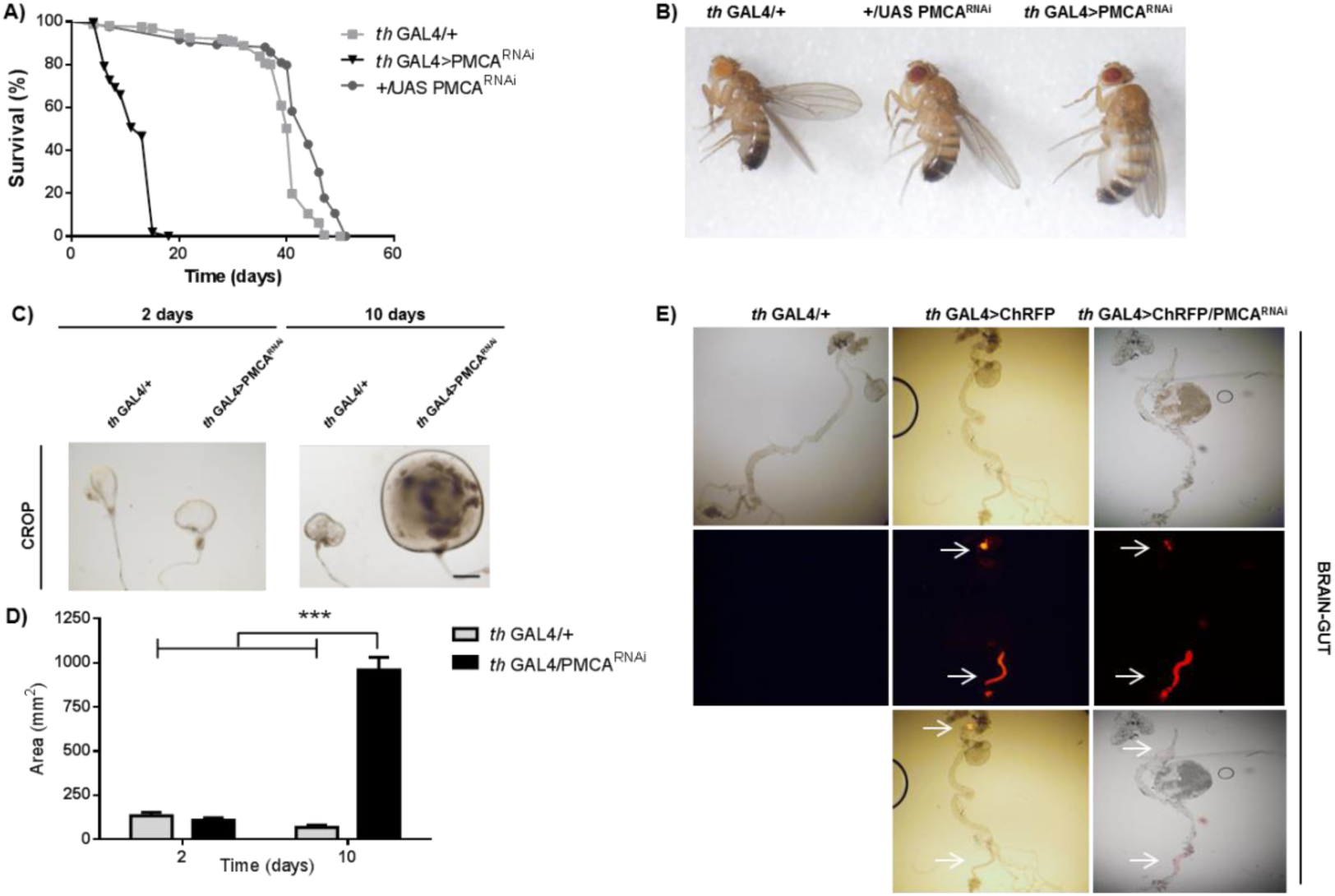
Flies th GAL4>PMCA^RNAi^ showed a shorter lifespan and digestive tract enlargement. A. *th* GAL4>PMCA^RNAi^ flies showed a shorter lifespan of ∼13 days. Flies from other genotypes died near 45 days. B, *Perimortem* swelling phenotype of *th* GAL4>PMCA^RNAi^ flies. C-D, Crop analyses. C, Photographs of crops from 2 and 10 days from flies expressing PMCA^RNAi^ in DN and control. D, Quantification of crop size. Flies expressing PMCA^RNAi^ in DN had larger crops than control only at 10 days after eclosion. E, Expression pattern of *th* driver (ChRFP) in gut. Light field images (upper), red fluorescence (middle) and merge (bottom). No expression was detected in *th* GAL4/+ gut (left); in contrast red signal was detected in proventriculus and hindgut (arrows) in *th* GAL4>ChRFP (middle) and *th* GAL4>ChRFP/PMCA^RNAi^ (right). C. Scale bar: 300 μm. A, Log-rank Mantel Cox test. D, Two-Way ANOVA Sidak test; ***p<0.001.

When we dissected the digestive tract, we observed an increment of crop size compared to controls (Figure 4C). Quantitative analyses showed 959,9±71,4 mm^2^ crops of *th-* GAL4>PMCA^RNAi^ vs. 68,7±11,7 mm^2^ of *th-*GAL4/+. This crop enlargement is significant only at 10 days post eclosion (Figure 4D). Therefore, effects of PMCA downregulation on survival and locomotor deficits could be dissociated since the latter occur not only at day 10 but also at day 2. Then, we wanted to know where in the digestive tract is the *th* driver active. Intestines of *th*-GAL4>ChRFP flies showed a strong red signal, principally in proventriculus and rectal ampulla (Figure 4E). Expression of PMCA^RNAi^ in proventriculus showed a lower GFP signal compared to control (Figure S3). The proventriculus is a structure at the junction of the foregut and midgut where the esophagus, midgut, and crop merge. It functions as a valve and regulates the passage of food into the anterior midgut and crop, which together perform the functions of the mammalian stomach [20]. Taking this into consideration, we thought that the crop enlargement could be generated by proventriculus dysfunction, consequence of PMCA^RNAi^ expression in this organ, and that could be regarded as a possible cause of death. Interestingly, constipation is a common non-motor feature in PD patients.

In summary, these data support that Ca^2+^ unbalance in DN impacts their physiology (ROS and dopamine increments) resulting in locomotor, lifespan and digestive alterations, some of them resembling features of PD.

## Materials and Methods

### Reagents

L-DOPA (Sigma).

### Drosophila strains

Fly stocks and crosses were raised on standard corn meal-yeast-agar medium under a 12 h/12 h light/dark cycle in 4-inch plastic vials. Flies were transferred to a fresh food medium every 2-4 days. Unless it says otherwise, crosses were done at 25°C and experiments made with 8-10 days old males maintained at 28°C (15 males per vial). *w*^1118^ #5905, *elav* ^*c155*^-GAL4 #458, *th* (*ple*)-GAL4 #8848, *gmr* GAL4 #1104 UAS mCD8:ChRFP #27391, UAS GCaMP6f #42747, UAS-cyto-Grx1-roGFP2 #67662, UAS-cyto-Orp1-roGFP2 [5]; UAS-mito-roGFP2-Grx1 #67664 and UAS-mito-roGFP2-Orp1 #67667 lines were obtained from Bloomington Stock Center; UAS PMCA RNAi #101743 was obtained from Vienna *Drosophila* RNAi Center.

### Ca^2+^ imaging

Flies were anesthetized with CO_2_ and brains were dissected in PBS and mounted in Mowiol (Calbiochem, San Diego, CA). Images were taken within 30 min after dissection using a Zeiss LSM 510 laser scanning confocal microscope. Samples were exited with an Argon laser at 488 nm and a helium-neon laser at 543 nm. Digital images were obtained at 20X focusing in dopaminergic axon network and mean intensity of a selected area value was quantified using ImageJ program.

### Reactive oxygen species quantification

Flies heads were fixed for 20 min at room temperature (RT) in 4% paraformaldehyde in PB. Dissected brains were mounted in Mowiol. Images were taken using a Zeiss LSM 880 Airyscan laser scanning confocal microscope. Probe fluorescence was excited sequentially at 405 and 488 nm (line by line) and detected at 500–530 nm and with a 543 nm laser. Digital images were obtained at 20X focusing in dopaminergic axon network and/or PPM3 dopaminergic neurons cluster. Fluorescence intensity of 405 nm (oxidized probe) and 488 nm (reduced probe) in a selected area was measured using ImageJ program and the ratio 405 nm/488 nm was made to quantify oxidation.

### Eye toxicity and Scanning Electron Microscopy

The electroscan was performed with a Scanning Electron Microscopy Quanta FEG 250, using low vacuum system and a Large Field Detector. The photomicrographs were taken at 5.00kV. To analyze toxicity in a quantitative manner, 7 eyes per genotype were randomly selected, orientated and recorded. The number of bristles per ommatidia in a constant area was counted using the point selection tool in ImageJ.

### Immunostaining

Flies’ heads were fixed for 40 min at RT in 4% paraformaldehyde in PB. Dissected brains were then blocked in 5% goat serum in phosphate-buffered saline with 0,1% Triton X-100 (PBS Tx) for 1h at RT and incubated with the primary antibody anti TH (*pale*) (1/50, Immunonstar) for 72 hs at 4 °C and then with the secondary antibody Cy5-conjugated donkey anti-mouse (1/1000, Jacksons Immunoresearch) for 2h at RT (brains were washed 3 times for 10 min with PBS Tx after each step) and mounted in Mowiol. Digital images were obtained at 20x and 40x with a Zeiss LSM 510 laser scanning confocal microscope. The dopaminergic neurons were counted manually through 40x Z-stacks. PPL1, PPL2, PPM1/2, PPM3 and PAL clusters were recorded from both hemispheres.

### Climbing assay (rapid iterative negative geotaxis, RING)

20 male flies were maintained per vial. Vertical mobility of 5- or 10-days old males was tested using the RING assay as described [12]. Briefly, the day before the assay 10 flies per genotype were randomly selected under CO_2_. The following day, each group was transferred into empty 4-inch glass vials without anesthesia and the vials were loaded into the RING apparatus. The apparatus was tapped three times in rapid succession to initiate a negative geotaxis response. Digital movies were taken and after 7 sec, the climbed distance in cm was measured for each fly and the average height per genotype calculated using the Scion Image software. Data from 5 technical replicates from each of 3 independent biological experiments per genotype were analyzed.

### Open field assay with video tracking

A panel of 8 arenas of 3.5 cm in diameter was used, allowing a simultaneous recording of 8 flies similar to [14] using a Logitech camera. Arenas were placed above an array of white LEDs. 2- or 10-days old males were cooled on ice for 3 min and placed one per plate. Flies were recorded at 8 am as they showed a peak of locomotor activity at that time. Movies of 10 min were analyzed by Fly tracker program designed by Bochicchio and Bodin (unpublished method).

### Locomotor behavior analysis

2 days-old adult males were placed in glass tubes with food and monitored for activity with infrared detectors and a computerized data collection system (Trikinetics) [21]. Activity was monitored in 12hs light/12hs dark for 10 days at 28°C similar to [22]. Collected data were analyzed by chi-square periodogram analysis employing the ClockLab software (Actimetrics, Wilmette, IL). A minimum of three independent experiments, including at least 15 flies that survived throughout the experiment, were analyzed per genotype.

### Dopamine determination

Flies were anesthetized with CO_2_ and brains were dissected in PBS and kept at −80 °C. Tissue was homogenized in 0.1 mL of 0.3 M perchloric acid, centrifuged for 15 min at 3000 g at 4 °C and the supernatant was frozen at −80 °C. Levels of dopamine were measured by high-pressure liquid chromatography coupled to electrochemical detection (HPLC-ED) using a Phenomenex Luna 5 lm, C18, 250 9 4.60 mm column (Phenomenex, Torrance, CA, USA) and an LC-4C electrochemical detector with glassy carbon electrode (BAS). The working electrode was set at +0.65 V vs. an Ag/AgCl reference electrode. The mobile phase contained 0.76 M NaH2-PO4-H2O, 0.5 mM EDTA, 1.2 mM 1-octane sulfonic acid and 5% acetonitrile; pH was adjusted to 3.0. Dopamine quantification was referred to total protein content. Proteins were measured using Pierce BCA Protein Assay kit (Thermo Fisher Scientific).

### Survival assay

Adults were collected during 24 h after imago hatching and a total of 84-268 males were monitored for survival. Experiments were performed in several replicates. The data are presented in the form of curve reflecting the percentage of changes in median lifespan between experimental and control variants Kaplan-Meier approach, according [23] with some modifications.

### Sweeling phenotype and gut dissection

Flies were anesthetized with CO_2_ and pictures were taken with a magnifying glass (Olympus MVX10 stereo microscope). To study brain-gut expressing *th* driver or crops flies were anesthetized with CO_2_ and organs were dissected in PBS and mounted in Mowiol. Digital images were obtained at 4x in digital Olympus BX60 microscope. Crops area was quantified using ImageJ. Proventriculi were fixed for 20 min at RT in 4% paraformaldehyde in PB and Hoechst staining for 10 min. Images were taken 40x using a Zeiss LSM 880 Airyscan laser scanning confocal microscope.

### RNA extraction and quantitative real time PCR analysis

For each experiment, 30-40 flies were collected and frozen for at least 1 h. Heads were mechanically isolated and total RNA of 30 heads per genotype was extracted using Trizol (Invitrogen Carlsbad, USA) according to the manufacturer’s protocol. An additional centrifugation step at 11,000 ×g for 10 min was used to remove the cuticles prior to the addition of chloroform. The concentration of total RNA purified from each sample was measured with a Nanodrop 2000 spectrophotometer (Thermo Scientific Walthman, MA). Retrotranscription of 0.5 µg of total RNA was performed using the Superscript II system (Invitrogen, Carlsbad, USA) with oligo (dT) (Thermo Fisher Scientific). PCR reactions were done using Platinum Taq Polymerase (Thermo Fischer Scientific) following the manufacturers’ instructions. Reactions were run in an Mx3005P Cycler (Stratagene, Santa Clara, CA). Data were analyzed using MxPRO-Mx3005P software. The primers used for PMCA amplification were F 5’ GCTCTTTTGGGATTTGTCCA’3; R 5’ ACTTCTCCAAAGTGCCCTCA; for RPL32, F 5’ ATGCTAAGCTGTCGCACAAATG; R GTTCGATCCGTAACCGATGT.

### Statistical analysis

Results are presented as mean ± SEM. Comparisons were performed using statistical test according experiments and it was detailed in each figure legends. Statistical significance level was p<0.05. Analyzes were done with GraphPad Prism 6.0.

## Author Contributions

The authors declare no competing interests.

## Acknowlegements

We thank to Mercedes Pianetti and Laboratorio de Microscopía Electrónica INTI – Mecánica; to Maria Jose Ferreiro and Rafael Cantera from Instituto Clemente Estable for their help with fly lines; to María Fernanda Ceriani and her group, and Pablo Wappner and his group from IIBBA-Leloir Institute for providing some fly lines and technical assistance. We thank to Andrés Hugo Rossi for microscope assistance. We also thank Hugo Adamo, Martin Perez and Susana Silberstein for discussions on experimental interpretation. This work was supported by grants ISN-CAEN 2015 and PIP485-CONICET to MCL, and PICT3336-ANPCyT to FJP and MCL. MCL, MSM, LF, CCF; CH, AC, EMC, FJP are members of the investigator career of CONICET. MIF is a technician from CONICET. BE, PAB are fellows of CONICET. BAS is support by a Foundation Barón fellowship.

