## Supplemental Figures for "“Plasma Membrane Calcium Atpase downregulation in dopaminergic neurons alters cellular physiology and behavior in Drosophila melanogaster”"

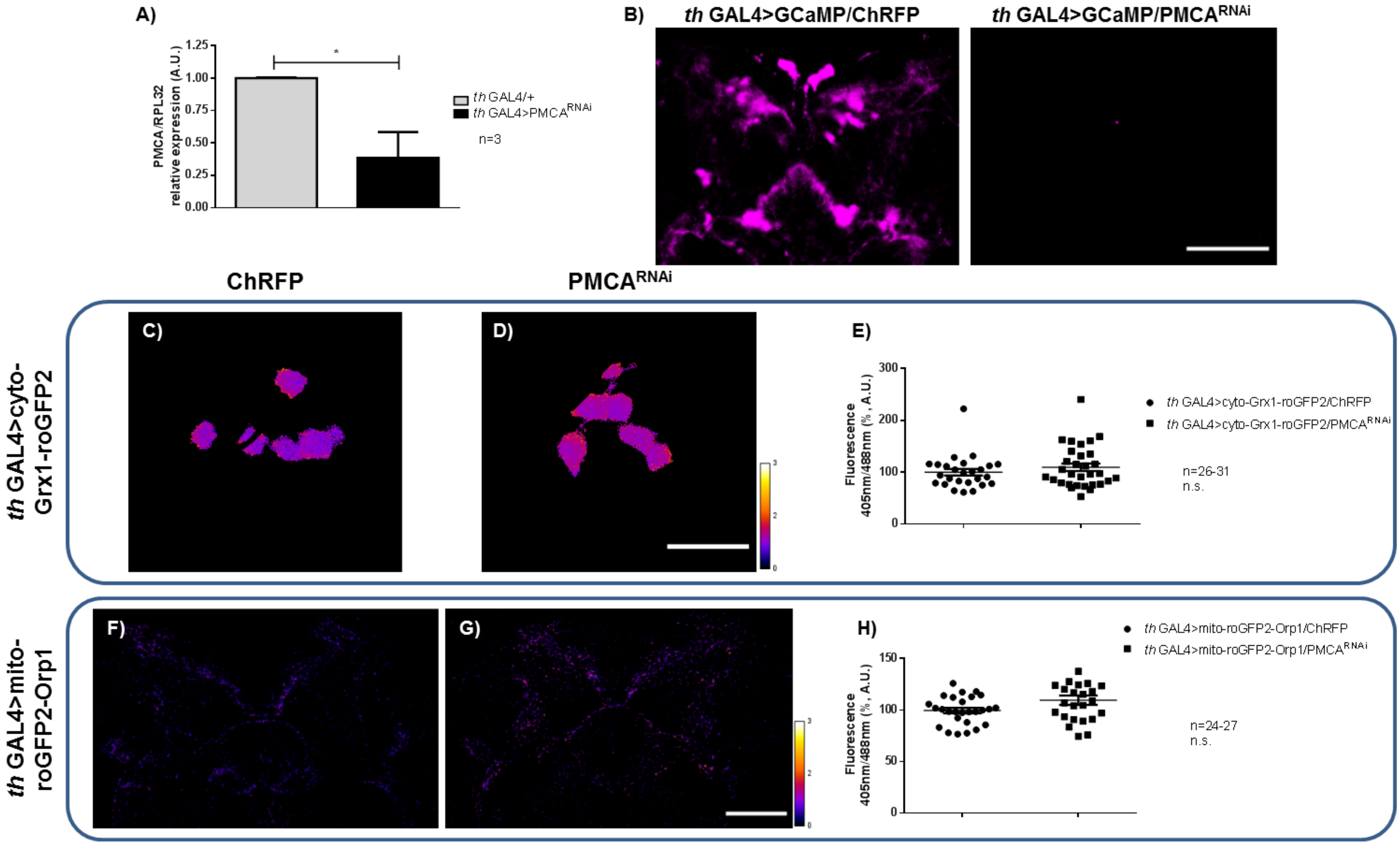


***Figure S1.*** A, Relative levels of PMCA mRNA measured by qPCR in heads. Expression of $PMCA^{RNAi}$ reduced PMCA mRNA ~40%. B, ChRFP signal (represented in magenta) detected in *th* GAL4>GCaMP/ChRFP axons (left); no signal was detected in *th* GAL4>GCaMP/$PMCA^{RNAi}$ (right). C-E, Photographs of PPM3 somas expressing C, ChRFP or D, $PMCA^{RNAi}$. E, Quantification 405nm/488nm fluorescence ratio of PPM3 cluster. No differences were found. F-H, Photographs of mitochondrial roGFP2-Orp1 in axons network of DN expressing F, ChRFP or G, $PMCA^{RNAi}$. H, Quantification of 405nm/488nm fluorescence ratio in each genotype. No differences were found. B. Scale bar: 100 μm. C, D. Scale bar: 15 μm. F, G. Scale bar: 50 μm. A, E, H, t test; n.s. p>0,05, *p<0.05.

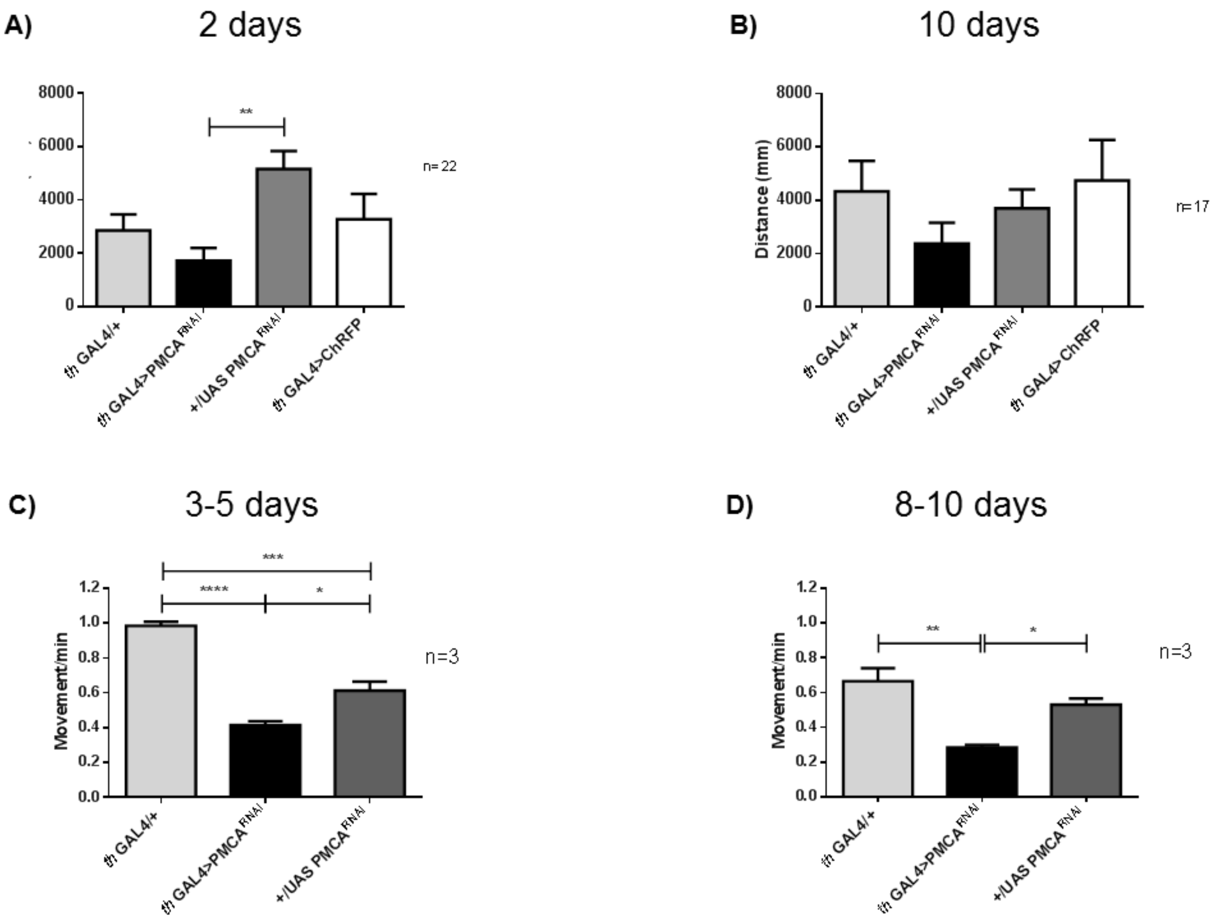


***Figure S2***. Locomotor activity. A, Total distance moved shows a significant reduction of movement only at 2 days post eclosion. B, No difference was observed at 10 days. C, Accumulated locomotor activity from 3-5 and 8-10 days after eclosion (D). Flies expressing $PMCA^{RNAi}$ moved less than controls in both cases. One-Way ANOVA, Tukey test; *p<0,05, **p<0,01 ***p<0,001, ****p<0,0001.

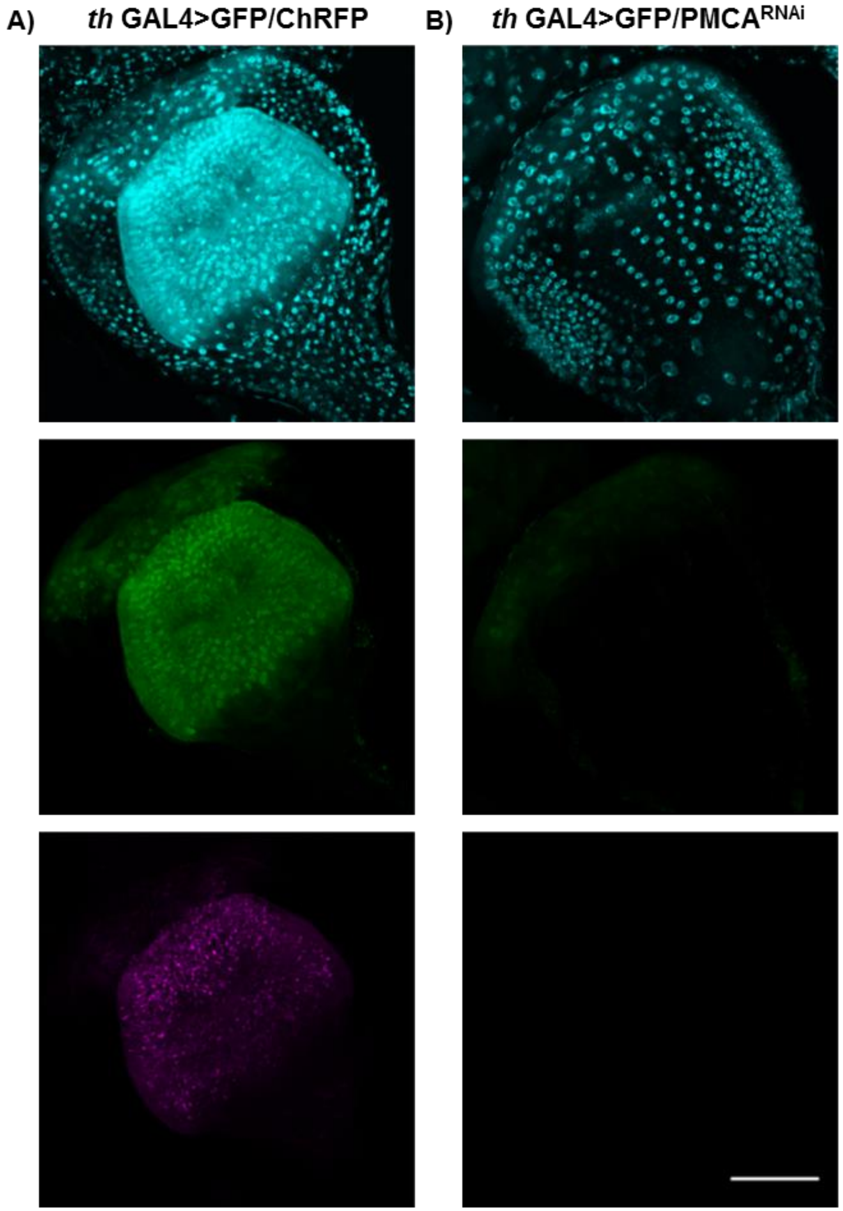


***Figure S3.*** Expression pattern of *th* driver in proventriculus of flies A, *th* GAL4>GFP/ChRFP (control) or B, *th* GAL4>GFP/PMCA$^{RNAi}$. Hoechst signal (upper), GFP signal (middle) and ChRFP signal (bottom, represented in magenta). GFP signal was reduced in PMCA$^{RNAi}$ expressing flies at 10 days after eclosion. A, B. Scale bar: 50 μm.
